# Development and Validation of a multi-target TaqMan qPCR method for detection of *Borrelia burgdorferi* sensu lato

**DOI:** 10.1101/2023.07.18.549442

**Authors:** Sébastien Masséglia, Magalie René-Martellet, Maxime Rates, Cecilia Hizo-Teufel, Volker Fingerle, Gabriele Margos, Xavier Bailly

## Abstract

Reliable detection of bacteria belonging to the *Borrelia burgdorferi* sensu lato species complex in vertebrate reservoirs, tick vectors, and patients is key to answer questions regarding Lyme borreliosis epidemiology. Nevertheless, the description of characteristics of qPCRs for the detection of *B. burgdorferi* s. l. are often limited. This study covers the development and validation of two duplex taqman qPCR assays used to target four markers on the chromosome of genospecies of *B. burgdorferi* s. l..

Analytical specificity was determined with a panel of spirochete strains. qPCR characteristics were specified using water or tick DNA spiked with controlled quantities of the targeted DNA sequences of *B. afzelii*, *B. burgdorferi* sensu stricto or *B. bavariensis*. The effectiveness of detection results was finally evaluated using DNA extracted from ticks and biopsies from mammals whose infectious status had been determined by other detection assays.

The developed qPCR assays allow exclusive detection of *B. burgdorferi* s. l. with the exception of the M16 marker which also detect relapsing fever *Borrelia* species. The limit of detection is between 10 and 40 copies per qPCR reaction depending on the sample type, the *B. burgdorferi* genospecies and the targeted marker. Detection tests performed on various kind of samples illustrated the accuracy and robustness of our qPCR assays.

Within the defined limits, this multi-target qPCR method allows a versatile detection of *B. burgdorferi* s. l., regardless of the genospecies and the sample material analyzed, with a sensitivity that would be compatible with most applications and a reproducibility of 100% under measurement conditions and limits of detection, thereby limiting result ambiguities.

**Highlights:** Four qPCR assays used in duplex were developed to detect *Borrelia burgdorferi* sensu lato. The limits of detection and quantification were defined according to state of the art standards. The specifications allow to detect *B. burgdorferi* sensu lato from different sampling sources.

## 1 Introduction

Lyme borreliosis (LB; also termed Lyme disease), the most common tick-borne disease in the temperate regions of the northern hemisphere, is caused by spirochetes of the *Borrelia (B.) burgdorferi* sensu lato (s. l.) complex. The bacteria are transmitted by vector ticks of the genus *Ixodes* (Stanek et al., 2012). Five *Borrelia* species are known to cause human Lyme disease in Europe and Asia: *B. burgdorferi* sensu stricto (s.s.) and *B. spielmanii* only in Europe, *B. afzelii, B. bavariensis* and *B. garinii* in Asia and Europe (Margos et al., 2019). Many vertebrates are blood-meal hosts for ticks and some of them are reservoirs of *Borrelia* spp.: small mammals for *B. burgdorferi* s.s., *B. afzelii*, *B. bavariensis*, and birds for *B. garinii* (Becker et al., 2016, Gern, 2008, Takano et al., 2011).

Detection methods of *B. burgdorferi* s.l. are applied on samples from: i) humans for disease diagnostic, ii) vertebrates to study pathogen ecology, iii) ticks, to measure the environmental risk of acquiring an infection (vector surveillance). Nowadays, several tests exist to detect *B. burdgorferi* s.l.. Indirect immunological approaches are more widely used for primary Lyme disease diagnostic in humans and animals due to their facility of implementation and standardization. Direct detection approaches, including DNA detection methods, can be used as complementary evidences to confirm infection and characterize the etiological agent, but it should not be used as screening test (Lager et al., 2017, Rauer et al., 2020, Schutzer et al., 2019).

Conversely, PCR or quantitative real-time PCR (qPCR), became widely used in *B. burgdorferi* s.l. ecology and vector surveillance (Rauter and Hartung, 2005, Strnad et al., 2020). TaqMan qPCR appears as a fast and reliable technology to study the transmission cycles of *B. burgdorferi* s.l. which involves analyzing a large sample number from different vertebrate hosts and ticks (Vourc’h et al., 2016). In the last two decades, many PCR and qPCR methods were developed to improve the sensitivity and specificity of the detection of *B. burgdorferi* s.l., including several commercial kits. The genes *recA*, *flaB*, *fliD*, *hbb*, 23S rRNA, 16S rRNA and p66 or the IGS 5S-23S located on the chromosome of strains of *B. burgdorferi* s.l. and the gene *ospA* located on the linear plasmid lp54 are commonly targeted (Courtney et al., 2004, Gooskens et al., 2006, Lager et al., 2017, Machackova et al., 2006, Ornstein and Barbour, 2006, Portnoi et al., 2006, Schwaiger et al., 2001, Strube et al., 2010, Tsao et al., 2004, Venczel et al., 2016).

Nevertheless, the description of the properties of detection methods and quantification for qPCR is often limited. Furthermore, the genomic diversity of *B. burgdorferi* s.l., at both the species and strains level, increases the difficulty to find a suitable detection test. While few published studies aim to compare or evaluate the specificity and sensitivity of TaqMan qPCR methods used to detect *B. burgdorferi* s.l. species and most of them for diagnostic purposes on human clinical samples (Briciu et al., 2016, de Leeuw et al., 2014, Ferdin et al., 2010, Maes et al., 2017, Nunes et al., 2018, Venczel et al., 2016), there is still a need for more standardized and characterized detection/quantification approaches.

In this study, we report the development of qPCR protocols targeting several loci for the detection and quantification of *B. burgdorferi* s.l.. We used available genomic sequences to design qPCR primers and probes that could be used to detect and quantify most species of *B. burgdorferi* s.l. from DNA extracted with or without tick DNA added. We verified using reference strains that the markers indeed worked properly for nine different species of the *B. burgdorferi* s.l. complex. We investigated the limits of detection and quantification of the markers for *B. afzelii*, *B. bavariensis* and *B. burgdorferi* s.s.. We finally questioned the efficiency of the developed qPCR protocols by studying the presence/absence of *B. burgdorferi* s.l. in samples previously characterized by other PCR based approaches.

## 2 Material and methods

### 2.1 Primers and probes

In order to design new protocols for the detection and quantification of *B. bugdorferi* s.l., we first developed new taqman qPCR assays targeting chromosomal loci of the bacterial species complex and another one to estimate the amount of tick DNA that could be used as a control for tick DNA extracts. The sequences of the designed oligonucleotides sequences are summarized in table 1.

**Table 1:**
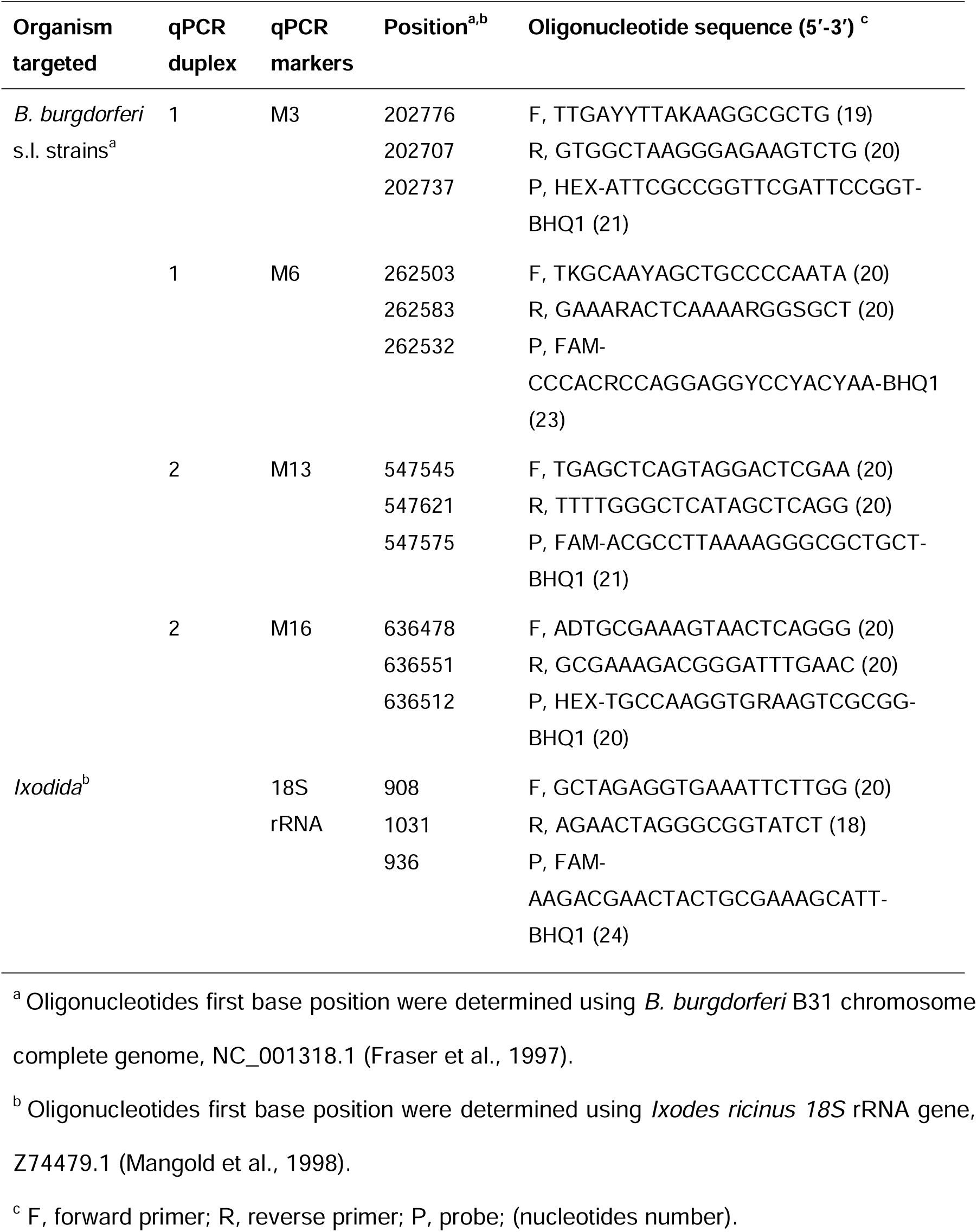
Primers and probes specially designed for this study.

Tick primers and a TaqMan probe were designed from 347 Ixodida *18S* rRNA sequences, including 17 genera and 111 species (Supplementary material 1). Sequences were aligned with the software Muscle (version 3.8.31) (Edgar, 2004), and oligonucleotides were designed to target conserved regions using the find primer tool of the software AliView (version 1.26) (Larsson, 2014). As explained above, we used these oligonucleotides targeting the *18S* rRNA gene of ticks as a control in parallel with *B. burgdorferi* s.l. detection (Camicas, 1998).

Eleven strains assigned to five species of the *B. burgdorferi* s.l. species complex that are publicly available through sequencing projects were chosen to represent the diversity of the targeted genomes based on available genomic datasets (Table 2).

**Table 2:** *B. burgdorferi* s.l. genomes used to design primers and probes.

| Organism | BioSample |
| --- | --- |
| *B. burgdorferi* sensu stricto N40 | SAMN14224735 |
| *B. burgdorferi* sensu stricto B31 | SAMN12121442 |
| *B. burgdorferi* sensu stricto 156a | SAMN12112595 |
| *B. burgdorferi* sensu stricto JD1 | SAMN09744444 |
| *B. bissettiae* DN127 | SAMN02604204 |
| *Borrelia afzelii* HLJ01 | SAMN02603021 |
| *Borrelia afzelii* PKo | SAMN02604205 |
| *Borrelia valaisiana* VS116 | SAMN12112918 |
| *Borrelia bavariensis* PBi | SAMN02603240 |
| *Borrelia bavariensis* BgVir | SAMN02603389 |
| *Borrelia bavariensis* NMJW1 | SAMN02603884 |

The chromosomes of these strains were obtained either from NCBI RefSeq corresponding accessions when the sequencing project was complete, or from *de novo* assemblies of raw reads obtained from the SRA archive using SPAdes 3.13.0 (Nurk et al., 2013).

Mauve 2.3.1 (Darling et al., 2010) was used to identify chromosomal sequences in *de novo* assemblies and to obtain a multiple-alignment of all the available chromosomes. Sequences of the different species are quite divergent. Therefore, we hypothesized that the ancestral sequence of the alignment would minimize the number of potential mismatches with all sequences of the *B. burgdorferi* s.l. species complex. Then, we inferred ancestral sequences at the internal nodes of a maximum likelihood tree based on the alignment of the studied chromosomes using the baseml program of the PAML 4.8 suite (Yang, 1997). The analysis was performed using a GTR model of sequence evolution with heterogeneous evolutionary rates modeled using a Gamma law discretized in five categories.

Primers and hybridizing oligos were selected in 48,000 kb windows along the whole chromosome of the most ancestral sequence using Primer-Blast (Ye et al., 2012). Parameter settings included: PCR product length between 70 and 100 pb, melting temperatures of the oligos between 55 and 60°C, and checking the specificity of primers and probes against *Ixodes* sp. sequences stored in the nr database. We identified polymorphisms that could affect the efficiency of the qPCR in priming site by blasting targeted loci, and introduced degeneracy in oligos based on the most problematic sites.

### 2.2 qPCR protocols

We first assessed the feasibility of amplification from seven qPCR targets, including tests of susceptibility to matrix inhibitors (data not shown), i.e. selectivity (Toohey-Kurth et al., 2020). We further tested two duplex qPCR to detect/quantify *B. burgdoferi* s.l. : one with primers and probes Marker 3 (M3) and Marker 6 (M6), another with Marker 13 (M13) and Marker 16 (M16). Taqman qPCR targeting the *18S* gene of ticks were performed independently to assess potential issues that could affect *B. burgdoferi* s.l. DNA detection in tick samples, e.g. low extraction yield or inhibitory effect. Taqman qPCR were performed in 96 well plates using a Bio-Rad CFX96 Real-time PCR thermocycler. Each reaction had a final volume of 25µl. It included 1x SsoAdvanced™ universal Probes Supermix (Bio-Rad, Hercules, USA), 200mM of each probe, 600mM of each primer. We added 5µl of DNA extract and, to reach the final reaction volume, molecular biology grade water potentially complemented with tick DNA. qPCR cycling conditions included a 3 min at 95°C initial step for polymerase activation and DNA denaturation, followed by 40 three-step cycles of i) denaturation at 95°C for 15 sec and ii) annealing-extension at 60°C for 30 sec and iii) fluorescence measurement.

The quantification cycle (Cq), determined using Bio-Rad CFX Manager, corresponds to the PCR cycles number when the fluorescent intensity increases above the baseline threshold set at 70 relative fluorescence units (RFU). A negative control of qPCR reaction mixes, containing 5µl of molecular biology grade water instead of the DNA extract was included on PCR plates.

### 2.3 Validation approach

The validation approach was based on the French standard NF U47-600 (AFNOR, 2015) which is congruent with the basic guide to real-time PCR in microbial diagnostics (Kralik and Ricchi, 2017) and the guidelines for validation of real-time PCR assays in veterinary diagnostic laboratories (Toohey-Kurth et al., 2020). Validation parameters were determined for the four markers using the duplex qPCR protocols described above. As described in figure 1, our validation protocol included: i) inclusivity and exclusivity tests based on cultured strains DNA; ii) determination of qPCR specifications based on synthetic targets DNA, iii) a study of detection results compared to published protocols using DNA of wild mammal’s and questing ticks.

#### 2.3.1 Inclusivity and exclusivity

We investigated the ability of the qPCR schemes to detect bacteria of the *B. burgdorferi* s.l. species complex and we identified the species that were exclusively detected using ten-fold dilution series of DNA extracts corresponding to 2000 to 0.02 spirochetes/µl obtained from cultures of fifteen *B. burgdorferi* s.l. strains, two relapsing fever *Borrelia* and two other spirochete strains (2000 spirochetes/µl) prepared by the German National Reference Center for *Borrelia*. Similar dilution series were previously used to compare real-time PCR protocols (panel III) (Lager et al., 2017). Two qPCR duplicates were performed for each of the 95 samples using both duplex Taqman assays targeting M3-M6 and M13-M16. The samples from which Cq are obtained for both duplicates are considered positive.

#### 2.3.2 qPCR specifications

Three plasmid inserts were designed using DNA sequences of three different strains respectively, 738bp of *B. burgdorferi* B31 (NRZ CP019767.1), 739bp of *B. bavariensis* NMJW1 (CP003866.1), 739bp of *B. afzelii* PKo (CP002933.1) to produce controlled amounts of standard qPCR targets. Each plasmid insert included the sequences of target M3, M16, M6, M13 cloned in an end-to-end fashion for a given reference to test the different qPCR using the same dilution series. These sequences were synthesized and inserted in pUC57 plasmids that carried a resistance gene to ampicillin. Sequences of plasmids were checked by Sanger sequencing (Genewiz, Leipzig, Germany). Plasmids were cloned in chemically competent cells of *Escherichia coli* DH5α using Subcloning Efficiency™ DH5α Competent Cells, Invitrogen™ (Thermo Fisher Scientific, Waltham, USA) and grown in LB broth medium with 100µg/ml ampicillin. Each standard plasmid was extracted from culture using NucleoSpin Plasmid (Macherey Nagel, Düren, Germany), quantified using a Qubit Fluorometer (Thermo Fisher Scientific, Waltham, USA), then diluted with sterile filtered water for molecular biology (Sigma-Aldrich, Saint-Quentin Fallavier, France) to produce an initial solution containing 10^9^ plasmids/µl and finally aliquots of 10^7^ plasmids/µl. Serial dilutions of plasmids were produced from these aliquots just before each qPCR plate preparation.

Two-fold dilution series from 40 to 1.25 plasmids per 5µl were prepared using sterile filtered water for molecular biology (Sigma-Aldrich, Saint-Quentin Fallavier, France). Eight repeats of the six dilutions were analyzed per plate. Three qPCR plate replicates were done, resulting in 24 qPCR reactions for each dilution. The qPCR limit of detection (LOD) was defined by the plasmid number that yielded a quantification cycle measure (Cq) for at least 95% of the 24 qPCR reaction repeats (AFNOR, 2015).

In order to detect a potential inhibitory effect of tick DNA that could impact sensitivity, we performed a similar experiment to assess the LOD by diluting each standard plasmid in DNA extracted from ticks free of *B. burgdorferi* s.l.. This tick DNA has been obtained from unengorged *I. ricinus* ticks reared in laboratory conditions (Leger et al., 2015). We targeted a concentration of tick DNA of 1.25ng/µl for the different samples, corresponding to a targeted Cq of 20. The qPCR targeting the 18S rRNA gene of *Ixodes* was used as internal control of the quantity homogeneity of tick DNA in every dilution. Two-fold dilution series of plasmids from 160 to 5 plasmids per 5µl were prepared. Four repeats of each dilution series were analyzed per plate. Two qPCR plate replicates were realized, resulting in eight qPCR reactions for each dilution. The qPCR LOD in tick DNA samples was defined as the plasmid number that yielded a Cq value for all of the eight qPCR reaction repeats. A negative qPCR control, containing 5µl of the non-infected tick DNA extract used as diluent was included on PCR plates.

The qPCR LOD in water and qPCR LOD in tick DNA solution were determined for the three standard plasmids *B. burgdorferi* B31_NRZ, *B. bavariensis* NMJW1, *B. afzelii* PKo using qPCR devices targeting M3-M6 and M13-M16.

To assess the linearity, amplification efficiency (E) and limit of quantification (LOQ), ten-fold dilution series of five logs, from 10^6^ to 10^2^ plasmids per 5µl, were prepared. Series of standard plasmid dilutions were analyzed using eight replicates in different qPCR plates.

At each *x_i_* level corresponding to the theoretical decimal logarithm (log10) of the plasmid number *x^′^_i_* we calculated : i) an observed value 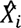 of the log10 plasmid number from the Cq value of each replicate using the equation of the calibration curve, ii) the mean 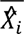 and the standard deviation *s*^′^ characterizing the fidelity from the 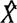 values of the eight replicates, iii) a coefficient of variation expressed in percentage using the formula 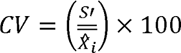, iv) a mean bias characterizing the precision of the experiment, measuring the gap between the average 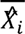 of observed values and the theoretical log10 plasmid number *x^′^_i_*, and v) an expanded linearity uncertainty value *U_LINi_* using the equation 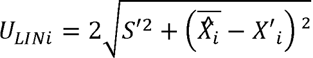.

We assessed the calibration curve linearity by evaluating whether the absolute value of the bias was less than 0.25*log_10_ at each dilution level (AFNOR, 2015).

The lowest plasmid quantity that showed a coefficient of variation under 25% was considered as the limit of quantification (LOQ) of the evaluated qPCR (Kralik and Ricchi, 2017). An expanded linearity uncertainty value for all of the *x_i_* levels, combining precision and fidelity, was calculated using the formula 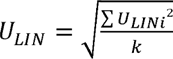. The mean of Cq values obtained for each dilution level were plotted against the theoretical log_10_ plasmid number *x*^’^ to obtain a linear calibration curve. The equation, the slope *a* and the regression coefficient *R*^²^ of the calibration curve were determined. The French standard NF U47-600 recommends to test series of at least four dilutions with eight replicates. Johnson et al. (2013) recommends a *R*^²^ higher than 0.98 over at least six logs and with three replicates. We analyzed eight series of five dilutions (AFNOR, 2015). qPCR efficiency was determined from each calibration curve using the equation 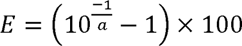. The qPCR efficiency should be between 90% and 105% (Johnson et al., 2013).

These analyzes were repeated for the three standard plasmids containing sequences representative of *B. burgdorferi*, *B. bavariensis* NMJW1 and *B. afzelii* Pko using the four markers amplified in duplex (M3-M6 and M13-M16).

#### 2.3.3 Detection results in various contexts

The sensitivity and specificity of both duplex qPCR schemes, targeting M3-M6 and M13-M16, were characterized with DNA samples from mammals and ticks for which detection of *B. burgdorferi* s.l. was previously reported using different methods: i) bank vole (*Myodes glareolus*) and chipmunk (*Tamia sibiricus*) DNA extracts analyzed using a PCR which targets the 16S rRNA gene confirmed using RFLP (Marconi and Garon, 1992, Postic et al., 1994); ii) DNA extracts of tick samples analyzed with a SYBR Green qPCR that targets the *flaB* gene, modified from Gómez-Díaz et al. (2010). DNA stored at −20°C had been extracted from mammalian ear biopsies using the NucleoSpin Tissue kit (Machery-Nagel, Düren, Germany) and from ticks using ammonia (Schouls et al., 1999). Samples that had previously shown an amplification were considered positive and others as negative. We used the developed duplex Taqman assays on: i) 47 positive and 76 negative bank voles samples, ii) 21 positive siberian chipmunks samples (Marsot et al., 2013), iii) 65 positive and 69 negative nymphs of *Ixodes ricinus* ticks samples (Vourc’h et al., 2016). To check the quality of the qPCR with a calibration curve, we analyzed plates containing each a 10-fold dilution series from 5*10^6^ to 5*10^2^ of standard plasmids. Positive samples identified positive with the new qPCR schemes were called Positive for the two assays: PP; those identified negative were called Positive initially then Negative: PN. Similarly, negative samples identified negative with the new qPCR schemes were called NN and those identified positive were called NP. The diagnostic sensitivity is equal to the percentage of PP for positive samples, and the diagnostic specificity is equal to the percentage of NN for negative samples (Kralik and Ricchi, 2017). In order to get further insights into the result of *B. bugdorferi* s.l. detection in tick DNA, the distributions of measured Cq with other tools and the newly developed primers were investigated further, both separately and in combination.

## 3 Results

### 3.1 Analytical sensitivity, limit of detection (LOD)

The limits of detection (LOD) by the new qPCR markers of *B. burdgorferi* s.s., *B. afzelii*, and *B. bavariensis* were first evaluated based on the number of standard plasmid copies diluted in water yielding a Cq value for 100% of the 24 replicates. LOD were similar for the three species: 40 copies per PCR reaction for the M3, and 20 copies per PCR reaction for the other markers: M6, M13 and M16 (Table 3). The positivity among the 24 repeats declined drastically from 100% to 0% below the observed LOD (Figure 2). The results of each replicate are detailed in supplementary material 2.

**Figure 1:**
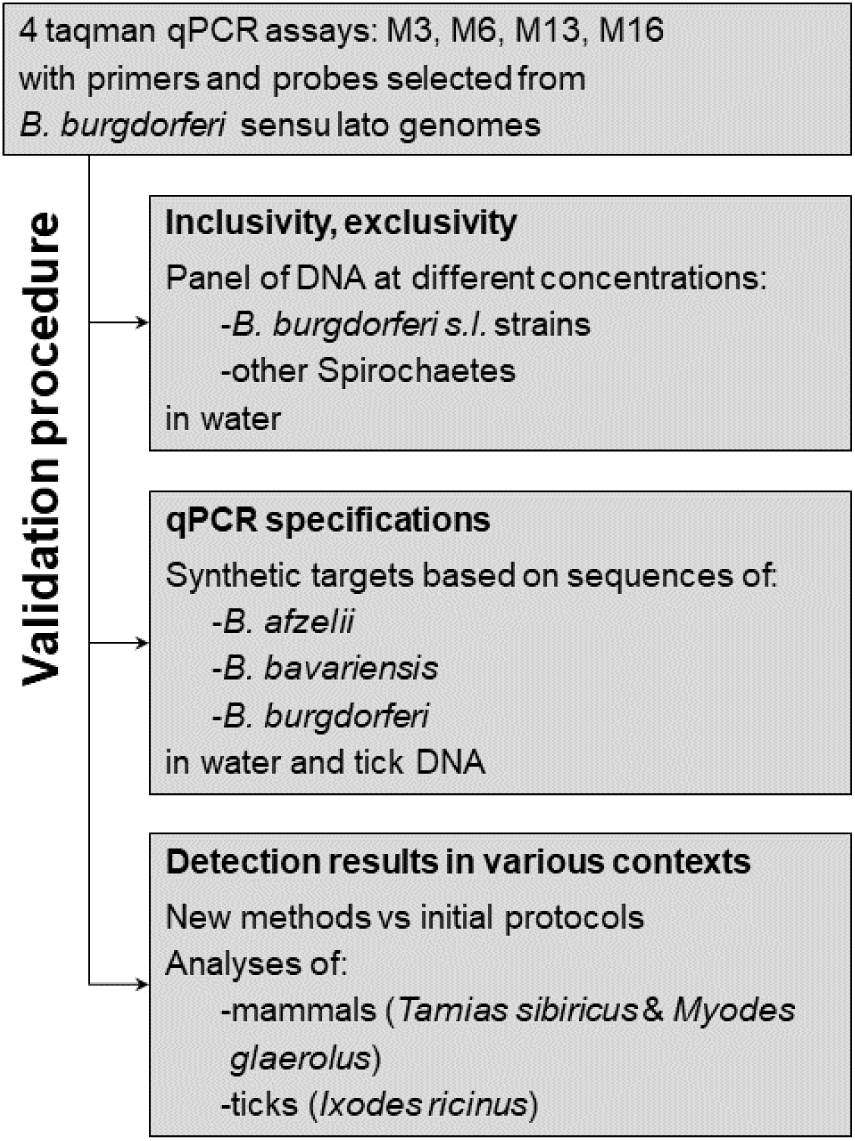
Summary of the three steps of the validation procedure.

**Figure 2:**
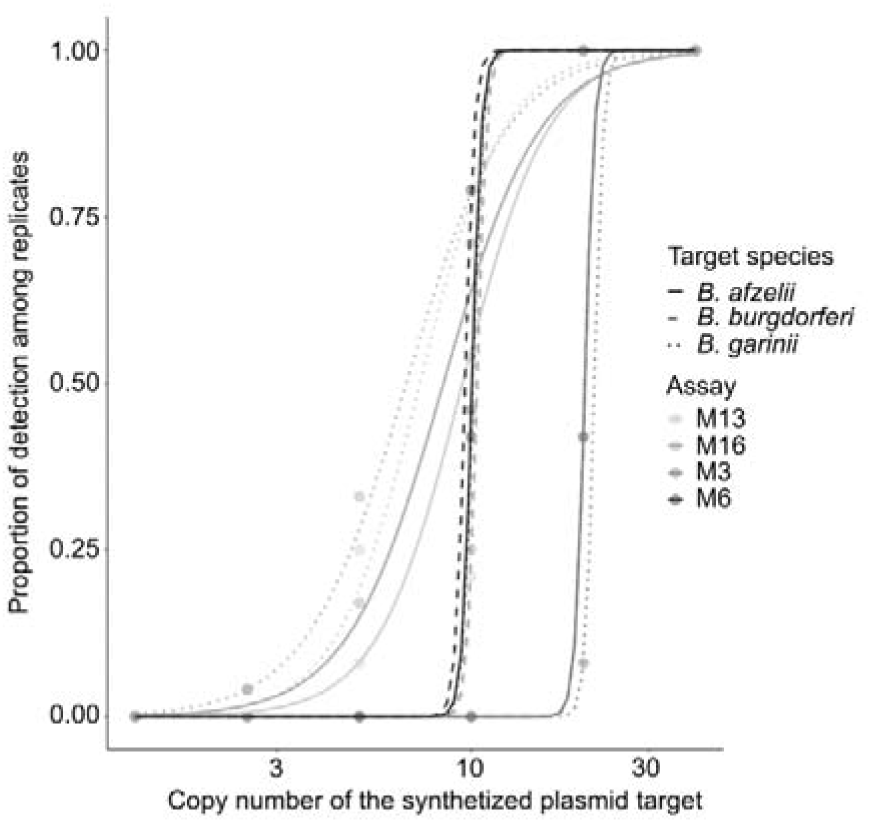
Lines describe the associated logistic relationship between the proportion of the 24 qPCR reaction repeats resulting in a detection for the four developed qPCR assays and the number of copies of the standard plasmids diluted in water for each marker and each targeted genospecies.

**Table 3:** Frequency of detection among 24 replicates of standard plasmids with inserts of the targeted sequences diluted in water for different *B. burgdorferi* s.l. species and different copy numbers. Mean of obtained Cq values is specified in brackets.

| Genome used<br>to design the<br>insert | Plasmid copy<br>number per<br>PCR reaction | In water without tick DNA |  |  |  |
| --- | --- | --- | --- | --- | --- |
|  |  | M3 | M6 | M13 | M16 |
| *B. afzelii* PKo | 40 | 100% (33.0) | 100% (31.3) | 100% (31.6) | 100% (32.3) |
|  | 20 | 42% (39.0) | 100% (35.3) | 100% (35.3) | 100% (35.9) |
|  | 10 | 0% | 46% (38.1) | 46% (38.7) | 54% (38.6) |
|  | 5 | 0% | 0% | 8% (38.7) | 17% (38.9) |
|  | 2.5 | 0% | 0% | 4% (38.7) | 4% (38.5) |
|  | 1.25 | 0% | 0% | 0% | 0% |
| *B. bavariensis*<br>NMJW1 | 40 | 100% (34.7) | 100% (31.3) | 100% (31.8) | 100% (32.5) |
|  | 20 | 8% (38.9) | 100% (34.9) | 100% (35.0) | 100% (35.6) |
|  | 10 | 0% | 42% (37.6) | 71% (38.5) | 71% (38.6) |
|  | 5 | 0% | 0% | 25% (38.9) | 33% (38.2) |
|  | 2.5 | 0% | 0% | 0% | 4% (39.6) |
|  | 1.25 | 0% | 0% | 0% | 0% |
| *B. burgdorferi*<br>B31_NRZ | 40 | 100% (33.3) | 100% (31.1) | 100% (31.8) | 100% (32.5) |
|  | 20 | 42% (39.2) | 100% (34.9) | 100% (35.5) | 100% (36.4) |
|  | 10 | 0% | 79% (38.5) | 21% (39.1) | 25% (38.9) |
|  | 5 | 0% | 0% | 0% | 0% |
|  | 2.5 | 0% | 0% | 0% | 0% |
|  | 1.25 | 0% | 0% | 0% | 0% |

The LOD was then evaluated using standard plasmids diluted in *I. ricinus* DNA free from *B. burgdorferi* s.l. DNA based on the number of copies yielding Cq values of 100% in eight replicates. We confirmed the lack of PCR inhibition and the homogeneity of the tick DNA concentration for all replicates, by obtaining a 19.8 mean Cq with a standard deviation of 0.1 cycles with the tick 18S rRNA gene qPCR. For M3, LOD were 20 copies per PCR reaction for the *B. afzelii* standard plasmid and 10 copies per PCR reaction for the *B. burgdorferi* s.l. and *B. bavariensis* standard plasmids. The LOD of the M6 was evaluated at 10 copies per PCR reaction of the three standard plasmids. For M13 and M16, LOD were evaluated at 10 copies per PCR reaction of *B. afzelii* standard plasmid and 5 copies per PCR reaction for the *B. burgdorferi* s.l. and *B. bavariensis* standard plasmids (Table 4). The results of each replicate are detailed in supplementary material 2.

**Table 4:** Frequency of detection among eight replicates of standard plasmids with inserts of the targeted sequences diluted in tick DNA for different *B. burgdorferi* s.l. species and different copy numbers. Mean of obtained Cq values is specified in brackets.

| Genome used to design the insert | Plasmid copy number per PCR reaction | With DNA of uninfected ticks |  |  |  |
| --- | --- | --- | --- | --- | --- |
|  |  | Duplex 1 |  | Duplex 2 |  |
|  |  | M3 | M6 | M13 | M16 |
| *B. afzelii* PKo | 160 | 100% (32.5) | 100% (31.4) | 100% (31.5) | 100% (32.2) |
|  | 80 | 100% (33.5) | 100% (32.6) | 100% (32.4) | 100% (33.2) |
|  | 40 | 100% (34.6) | 100% (33.5) | 100% (33.8) | 100% (34.6) |
|  | 20 | 100% (35.9) | 100% (34.5) | 100% (34.2) | 100% (35.0) |
|  | 10 | 87% (36.2) | 100% (35.3) | 100% (35.4) | 100% (36.2) |
|  | 5 | 87% (37.8) | 87% (36.0) | 87% (36.0) | 87% (37.0) |
| *B. bavariensis* NMJW1 | 160 | 100% (32.8) | 100% (30.3) | 100% (30.3) | 100% (31.1) |
|  | 80 | 100% (33.8) | 100% (31.3) | 100% (31.3) | 100% (32.0) |
|  | 40 | 100% (34.6) | 100% (32.2) | 100% (32.6) | 100% (33.3) |
|  | 20 | 100% (36.2) | 100% (33.4) | 100% (33.6) | 100% (34.3) |
|  | 10 | 100% (37.3) | 100% (34.4) | 100% (34.5) | 100% (35.3) |
|  | 5 | 62% (38.4) | 87% (36.0) | 100% (35.9) | 100% (37.1) |
| *B. burgdorferi* B31_NRZ | 160 | 100% (32.6) | 100% (30.6) | 100% (31.0) | 100% (31.6) |
|  | 80 | 100% (33.8) | 100% (31.8) | 100% (32.0) | 100% (32.7) |
|  | 40 | 100% (34.7) | 100% (32.8) | 100% (32.9) | 100% (33.6) |
|  | 20 | 100% (35.8) | 100% (33.8) | 100% (34.0) | 100% (34.8) |
|  | 10 | 100% (36.8) | 100% (34.9) | 100% (35.0) | 100% (36.0) |
|  | 5 | 75% (38.4) | 75% (36.1) | 100% (36.0) | 100% (37.0) |

Taking into account the results of the different plasmids, the evaluated LOD of *B. burgdorferi s.s.*, *B. afzelii*, and *B. bavariensis* in infected *I. ricinus* DNA extracts was at 20 copies per PCR reaction for M3 and 10 copies per PCR reaction for M6, M13 and M16.

### 3.2 Limit of quantification (LOQ)

The regression coefficient *R*^²^ of all replicates of all calibration curves was above the required minimum value of 0.98. Depending on the standard plasmid, the qPCR efficiency was between 97% and 101% for M3 and between 96% and 100% for M6 for the first duplex qPCR scheme. For the second duplex qPCR scheme, efficiencies varied between 98% and 102% for M13 and between 99% and 103% for M16. The absolute bias value was under 1.2 copies and the coefficient of variation was under 2.7% for all markers, the three standard plasmids and the different levels of the dilution range (between 10^6^ to 10^2^ plasmids). Based on thresholds of 0.25 log10 at each dilution level for the absolute bias value and 25% for the coefficient of variation at the LOQ, the linearity of the calibration curve is validated and the LOQ fixed at 100 copies of standard plasmid in the 5μl of DNA extract used as template. The expanded linearity uncertainty value of both qPCR devices for all dilution levels was between 1.1 and 1.3 copies (Table 5). The results of each replicate are detailed in supplementary material 3.

**Table 5:** Parameters of linearity, CV with 10^2^ plasmids, expanded linearity uncertainty value (*U_LIN_*), R-squared (*R*^²^) and amplification efficiency (*E*) calculated from calibration curves of qPCR devices targeting M3-M6 and M13-M16

| Markers | Standard plasmid designed from | CV at LOQ | $U_{LIN}$<br>log<sub>10</sub> | $R^2$ | slope $a$ | $E$ |
| --- | --- | --- | --- | --- | --- | --- |
| M3 | *B. afzelii* PKo | 1.877% | 0.0499 | 0.9989 | -3.396 | 97% |
|  | *B. bavariensis* NMJW1 | 1.156% | 0.0784 | 0.9960 | -3.300 | 101% |
|  | *B. burgdorferi* B31_NRZ | 1.832% | 0.1009 | 0.9978 | -3.306 | 101% |
| M6 | *B. afzelii* PKo | 1.515% | 0.0628 | 0.9984 | -3.415 | 96% |
|  | *B. bavariensis* NMJW1 | 1.779% | 0.0912 | 0.9937 | -3.343 | 99% |
|  | *B. burgdorferi* B31_NRZ | 2.064% | 0.0915 | 0.9972 | -3.318 | 100% |
| M13 | *B. afzelii* PKo | 1.736% | 0.0522 | 0.9988 | -3.361 | 98% |
|  | *B. bavariensis* NMJW1 | 2.387% | 0.0792 | 0.9962 | -3.271 | 102% |
|  | *B. burgdorferi* B31_NRZ | 2.364% | 0.1049 | 0.9964 | -3.292 | 101% |
| M16 | *B. afzelii* PKo | 2.175% | 0.0542 | 0.9988 | -3.355 | 99% |
|  | *B. bavariensis* NMJW1 | 2.455% | 0.0846 | 0.9960 | -3.260 | 103% |
|  | *B. burgdorferi* B31_NRZ | 2.654% | 0.1025 | 0.9968 | -3.283 | 102% |

### 3.3 Analytical specificity

Negative results were obtained using a template DNA extract from 10,000 *Leptospira* and *Treponema* spirochetes. Similarly, targeting M3, M6 and M13 using 10,000 spirochetes of the relapsing fever species *Borrelia miyamotoi* and *Borrelia hermsii* resulted in negative results (Table 6). Nevertheless, targeting the M16, positive results were obtained from 10,000 of these relapsing fever spirochetes which demonstrates that M16 was amplified in a *Borrelia* species which does not belong to the group *B. burgdorferi* s.l.

**Table 6.** Results from the panel III of the German National Reference for *Borrelia* consisting of DNA extracted from 15 *B. burgdorferi* s. l. strains (10 fold dilution series corresponding to 2000 to 0.02 spirochetes/µl), two relapsing fever *Borrelia* and two other spirochete strains (Leptospira, Treponema) (Lager et al., 2017). The lowest spirochete number with positive result is specified with Cq values in brackets.

| Strains | Spirochete number per reaction (Cq) |  |  |  |
| --- | --- | --- | --- | --- |
|  | Duplex 1 |  | Duplex 2 |  |
|  | M3 | M6 | M13 | M16 |
| *B. burgdorferi* s.s. B31 | 10 (34.7 - 34.1) | 1 (38.5 - 35.8) | 10 (31.8 - 32.7) | 10 (33.4 - 32.9) |
| *B. burgdorferi* s.s. PBre | 10 (34.5 - 34.1) | 10 (31.9 - 31.5) | 10 (31.9 - 32.2) | 10 (32.3 - 32.3) |
| *B. afzelii* PKo | 10 (33.3 - 33.2) | 10 (31.4 - 31.3) | 1 (38.9 - 36.7) | 1 (39.9 - 39.6) |
| *B. afzelii* PVPM | 10 (33.7 - 33.7) | 10 (31.2 - 31.8) | 10 (31.6 - 31.6) | 10 (32.7 - 32.2) |
| *B. garinii* PBr | 10 (39.6 - 35.2) | 1 (38.6 - 37.8) | 10 (32.4 - 32.1) | 10 (32.1 - 33.6) |
| *B. garinii* PHei | 10 (36.1 - 37.0) | 10 (34.0 - 33.6) | 1 (39.2 - 39.5) | 10 (34.4 - 33.7) |
| *B. garinii* PWudII | 10 (34.2 - 34.5) | 10 (32.8 - 32.5) | 10 (31.7 - 31.9) | 10 (33.2 - 32.8) |
| *B. garinii* PRef | 10 (35.3 - 34.5) | 10 (32.7 - 32.5) | 10 (33.4 - 34.2) | 10 (34.4 - 33.5) |
| *B. garinii* PLa | 10 (35.1 - 36.8) | 10 (33.9 - 33.5) | 10 (34.0 - 33.5) | 10 (36.0 - 35.9) |
| *B. spielmanii* PSigII | 10 (36.0 - 39.0) | 100 (36.1 - 35.9) | 10 (33.0 - 33.6) | 10 (33.1 - 34.5) |
| *B. bavariensis* PBi | 10 (35.6 - 36.0) | 10 (34.3 - 34.4) | 10 (34.8 - 33.4) | 10 (33.6 - 36.8) |
| *B. bissetii* PGeb | 10 (35.6 - 36.0) | 100 (33.7 - 34.2) | 10 (31.8 - 31.2) | 10 (32.3 - 32.1) |
| *B. lusitanae* Poti B2 | 1000 (34.6 - 34.5) | 1 (37.0 - 38.7) | 10 (31.6 - 31.9) | 1 (37.7 - 36.1) |
| *B. valaisiana* VS116 | 10 (37.1 - 38.2) | 10 (32.8 - 32.6) | 10 (33.0 - 32.6) | 10 (33.6 - 33.5) |
| Reptile-associated *Borrelia* sp. | 1000 (34.2 - 34.6) | none | 10 (35.6 - 33.4) | 10 (36.1 - 37.9) |
| *B. hermsii* | none | none | none | 10000 (26.0 - 26.1) |
| *B. miyamotoi* | none | none | none | 10000 (22.8 - 22.9) |
| *Treponema phagedenis* | none | none | none | none |
| *Leptospira* | none | none | none | none |

Regardless of the targeted marker, positive results were found down to 10 spirochetes per PCR reaction for the strains *B. burgdorferi* s.s. PBre, *B. afzelii* PVPM, *B. garinii* PWudII, *B. garinii* PRef, *B. garinii* PLa, *B. bavariensis* PBi and *B. valaisiana* VS116. For the strains *B. burgdorferi* s.s. B31 and *B. garinii* PBr, positive results were found down to 1 spirochete per PCR reaction by targeting M6 and down to 10 spirochetes per PCR reaction by targeting all other markers. For *B. garinii* PHei, positive results were found down to 1 spirochete per PCR reaction by targeting M13 and down to 10 spirochetes per PCR reaction by targeting the other markers. For *B. afzelii* PKo, positive results were found for the two targeted markers down to 1 spirochete per PCR reaction using M3 and M13 and down to 10 spirochetes per PCR reaction for M6 and M16. Positive results were obtained for *B. lusitaniae* PotiB2 down to 1 spirochete per PCR reaction for M6 and M16, 10 spirochetes per PCR reaction for M13 and 1000 spirochetes per PCR reaction for M3. For the strains *B. spielmanii* PSigII and *B. bissettiae* PGeb, positive results were found down to 100 spirochetes per PCR reaction by targeting the M6 and down to 10 spirochetes per PCR reaction by targeting all other markers.

For reptile-associated *Borrelia sp.*, positive results were found down to 10 spirochetes per PCR reaction for the two markers targeted by the duplex qPCR scheme M13-M16, the other duplex PCR scheme only allow to detect more than 1000 bacteria per PCR reaction for M3 and did not yield positive results for M6.

### 3.4 Diagnostic sensitivity and specificity

#### 3.4.1 Myodes glareolus samples

Using the M3, M13 and M16 of the two duplex qPCR schemes, we detected a significant amplification signal from the 47 bank voles previously identified as infected by *B. burgdorferi s.l.* (diagnostic sensitivity = 100%). However, only two positive results were obtained using the M6 (diagnostic sensitivity = 4.3%). All 76 bank voles DNA extracts previously classified free of *B. burgdorferi* s.l. infection tested negative with M3 and M6 (diagnostic specificity = 100%). From the same 76 DNA extracts, one positive result was obtained targeting M13 and M16 (diagnostic specificity = 98.7%).

#### 3.4.2 Tamias sibiricus samples

Positive qPCR results were obtained using the developed duplex schemes from all 21 DNA extracts from *Tamia sibiricus* previously identify as infected by *B. burgdorferi* s.l., corresponding to a 100% diagnostic sensitivity for the four markers. The diagnostic specificity could not be evaluated because no DNA extracted from uninfected Siberian chipmunks was analyzed.

#### 3.4.3 Ixodes ricinus samples

From the 72 tick DNA extracts previously classified negative, i.e. not infected by *B. burgdorferi* s.l., one sample was detected positive using M3 and M6 of the first duplex qPCR scheme (diagnostic specificity = 98.6%), two samples were tested positive using M13 (diagnostic specificity = 97.2%) and none tested positive targeting M16 (diagnostic specificity = 100%). From the 65 tick DNA extracts previously classified positive, i.e. infected by *B. burgdorferi* s.l. (Vourc’h et al., 2016), only 34 DNA extracts presented a positive result targeting M3 (diagnostic sensitivity = 52.3%), 35 DNA extracts targeting M6 (diagnostic sensitivity = 53.8%), 36 DNA extracts present a positive result targeting M13 (diagnostic sensitivity = 55.4%) and 38 DNA extracts targeting M16 (diagnostic sensitivity = 58.5%).

Driven by the low diagnostic sensitivity observed for the tick DNA extracts previously identified as positive for *B. burgdorferi* s.l., we investigated the distribution of initial Cq values obtained previously with a SYBR Green qPCR assay targeting *flaB* gene modified from Gómez-Díaz et al. (2010) and those obtained with the new taqman procedures (Figure 3). The observed results illustrated two distinct patterns (Figure 3A): i) for Cq values of the published SYBR Green qPCR assay below 35 (Vourc’h et al., 2016), most of the tests involving the developed qPCR resulted in a Cq value, i.e. the detection of the pathogen. Among 30 tick DNA extracts initially found infected, 29 (diagnostic sensitivity = 96.7%), 28 (diagnostic sensitivity = 93.3%), 30 (diagnostic sensitivity = 100%) and 30 (diagnostic sensitivity = 100%) samples were detected positive using M3, M6, M13 and M16 respectively; ii) for Cq values of the published SYBR Green qPCR assay above 35, most of the tests based on the developed qPCR did not yield a positive amplification. Among 35 tick DNA extracts, only 5 (diagnostic sensitivity = 14.3%), 7 (diagnostic sensitivity = 20.0%), 6 (diagnostic sensitivity = 17.1%), and 8 (diagnostic sensitivity = 22.8%) DNA extracts resulted in a Cq value for M3, M6, M13 and M16, respectively. Based on the possible comparisons, there is a strong agreement between the initial Cq values and those obtained with the developed markers (Figure 3B).

**Figure 3:**
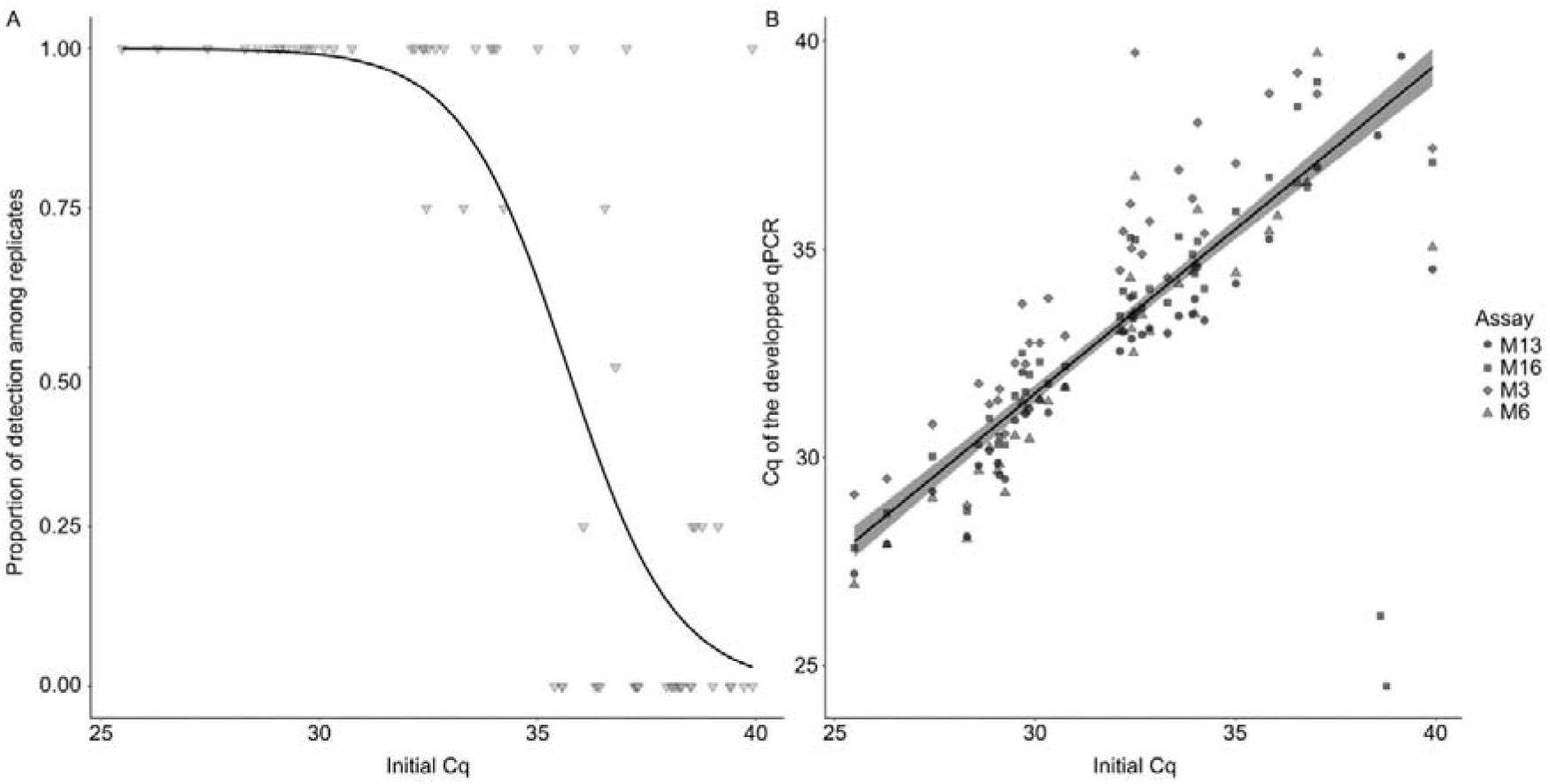
Evaluation of the results of real-time quantitative qPCR detection of *B. burgdorferi* s.l. developed Taqman assays based on tick samples previously evaluated as positive using another, non-gold-standard, SYBR Green method. A: Individual measures and logistic relationship between the proportion of the four developed qPCR assays resulting in a detection for a given tick and the Cq evaluated with the SYBR Green method used previously. B: Inidividual measures and linear relationship between initial Cq obtained from the SYBR Green method and Cq obtained from the developed assays.

## 4 Discussion

*Borrelia burgdorferi* s.l. is a species complex of vector borne bacterial pathogens. Some genotypes of this species complex are responsible of human Lyme borreliosis, a significant public health problem. In order to confirm medical diagnostic, to monitor the bacteria in vectors, or to study the bacterial ecology, there is a need to develop fully characterized tools for the detection and quantification of the bacteria in different contexts and for different applications. Here, we putting forward four taqman qPCR assay which can be used to detect and quantify *B. burgdorferi* s.l..

We first analyzed the efficiency of the different markers considering the diversity of *B. burgdorferi* s.l. based on the analysis of standard plasmids containing the homologous sequence of targeted loci for *B. burgdorferi* s.s., *B. afzelii*, and *B. bavariensis.* The efficiency and the linearity of the different qPCR assays used in duplex meet the quality criteria defined according to state of the art of qPCR characterization. For the three species targeted, detection is possible at 95% in 5µl of water spiked with less than 40 standard plasmids targeting M3 and less than 20 standard plasmids targeting M6, M13 and M16. We have tested that the quantification is reliable down to 100 target copies for all the markers for *B. burgdorferi* s.s., *B. afzelii*, and *B. bavariensis*. Limits of detection of plasmids can decrease to 20 copies in a solution of tick DNA targeting M3 and less than 10 standard plasmids targeting M6, M13 and M16. It is noteworthy that if it is possible to obtain Cq value from a spirochete number below the LOD or obtaining a quantity below the LOQ, the reliability and the repeatability of these results must be put into perspective (Kralik and Ricchi, 2017). In order to increase further the sensitivity of detection and quantification, the qPCR assay could be used in a digital droplet PCR framework, which give an absolute quantification of *Borrelia* (King et al., 2017).

We then analyzed the efficiency of the different markers considering the diversity of *B. burgdorferi* s.l. based on the analysis of DNA obtained from representative isolates of different species of the *B. burgdorferi* s.l. species complex as well as representatives of other spirochete species. DNA from the strain panel allowed the characterization of the analytical specificity of duplex qPCR assays M3-M6 and M13-M16. Regarding specificity, spirochetes not belonging to the genus *Borrelia* could not be detected. Potential detection of *B. hermsii* and *B. miyamotoi* that do not belong to the *B. burgdorferi* s.l. complex was observed using M16 only. The study of the spirochete strain panel allowed characterization of the analytical specificity of qPCR assays in a duplex setting. To this end, Cq values were obtained for all four targeted markers from all spirochetes belonging to the *B. burgdorferi* s.l. complex. Our results illustrate that having several alternative assays to detect pathogen genotypes can be useful, especially in settings where relapsing fever and Lyme *Borrelia* species overlap as is the case with *B. miyamotoi*. Since epidemiological or ecological studies usually use host or vector tissue to investigate the presence of *Borrelia*, we tested the specificity and sensitivity of the developed assays in *Ixodes* vectors and different hosts. This enabled us to compare the results they yielded to the results obtained with other published detection methods. With regard to the detection of *B. burgdorferi* s.l. in hosts, we identified a strong agreement of the developed markers with the previously initially used detection method. All samples from hosts initially detected positive were found positive. We just found one positive samples on 76 *M. glareolus* initially detected negative. Nevertheless, the experiments revealed that M6 could not be used to detect and quantify *B. burgdorferi* s.l. in bank voles. Among other reasons, this might be due to an unspecific hybridization of the M6 primers or probe on *M. glaerolus* DNA. The qPCR tests for *B. burgdorferi* s.l. are used on many different matrices (Estrada-Pena et al., 2016, Ivanova et al., 2014, Margos et al., 2011, Munoz-Leal et al., 2020). While it is not possible to systematically validate the sensitivity and specificity of the assays for each matrix, our results illustrate that it is critical to use multiple qPCR targets, well-characterized in other contexts, to ensure the quality of the results. Regarding the detection of *B. burgdorferi* s.l. in ticks, the comparison of the detection of *B. burgdorferi* s.l. with our taqman assays and previously used protocols suggests that the markers we have designed and tested in this study provide robust results : i) measured Cq values obtained from the developed qPCR schemes and previously used assay showed a strong correlation, ii) for samples that were found infected using a SYBR Green assay with a Cq value less than 35 (Vourc’h et al., 2016) our new qPCR schemes share similar sensibility properties to detect *B. burgdorferi* s.l.. However, for samples that were found infected using a SYBR Green assay with a Cq value above 35 (Vourc’h et al., 2016) our new qPCR is ineffective for more than 80% of them. This could for instance be explained either by a higher limit of detection or a higher specificity of our four qPCR assays compared to the formerly used SYBR Green assay or degradation of the DNA of the analyzed extracts. The properties of the SYBR Green assay have not been properly evaluated and it should not be taken as a gold standard. Our new assays would detect *B. burgdorferi* s.l. in 100% of cases from an infected *Ixodes ricinus* tick with a minimum load of 100 spirochetes. From a biological point of view, the usual bacterial load in ticks, ranged from 7969 to 84841 spirochetes per *I. ricinus* nymph (Herrmann et al., 2013). Our low LOD, imply that the assays we developed would provide a conservative and robust result. Overall, we developed four qPCRs assays to detect and quantify *B. burgdorferi* s.l.. qPCR targeting M13 has all the expected properties (Toohey-Kurth et al., 2020): i) It gave relevant results for the different DNA sources we tested ii) good repeatability, iii) good analytical specificity including inclusivity, exclusivity, and thus selectivity, iiii) good diagnostic sensitivity and diagnostic specificity. Nevertheless, taking into account the wide variety of bacterial genospecies targeted and the large number of hosts and vector that could be infected, we advocate that the four markers can complement each other to obtain more robust multiplex qPCR protocols. For instance, we validated two duplex assays M3 with M6 and M13 with M16.

## 6 CRediT authorship contribution statement

**Sébastien Masséglia:** Conceptualization, Methodology, Investigation, Validation, Writing - Original Draft, Writing - Review & Editing. **Magalie René-Martellet:** Conceptualization, Writing - Review & Editing, Supervision. **Maxime Rates:** Writing - Review & Editing. **Cecilia Hizo-Teufel:** Resources, Writing - Review & Editing. **Volker Fingerle:** Conceptualization, Resources, Validation, Writing - Review & Editing. **Gabriele Margos:** Conceptualization, Resources, Validation, Writing - Review & Editing. **Xavier Bailly:** Conceptualization, Formal analysis, Validation, Writing - Review & Editing, Supervision, Project administration.

## 7 Acknowledgement

The authors would like to thank Sarah Bonnet (UMR BIPAR ENVA-ANSES-INRAE, Maisons-Alfort, France) and Karen McCoy (UMR MIVEGEC CNRS-IRD, Montpellier, France) for providing the DNA extracted from ticks, reared in laboratory conditions, free from *B. burgdorferi* s.l. infection. We thank the « Tiques et Maladies à Tiques » (TMT) group of the « Réseau Ecologie des Interactions Durables » (REID) for discussion and support.

## 8 Supplementary data

Supplementary material 1 table of 18S rRNA sequences used to design the tick primers and TaqMan probe

Supplementary material 2 table of all limit of detection data

Supplementary material 3 table of all limit of quantification data

## 9 Data availability

Data will be made available on request.

## References

AFNOR, 2015. Animal health analysis methods - PCR - Part 2: requirements and recommendations for the development and the validation of veterinary PCR - NF U47-600-2 AFNOR Editions, pp. 51.

Becker, N.S., Margos, G., Blum, H., Krebs, S., Graf, A., Lane, R.S., Castillo-Ramirez, S., Sing, A., Fingerle, V., 2016. Recurrent evolution of host and vector association in bacteria of the *Borrelia burgdorferi* sensu lato species complex. BMC Genomics. 17, 734. https://doi.org/10.1186/s12864-016-3016-4.

Briciu, V.T., Sebah, D., Coroiu, G., Lupse, M., Carstina, D., Tatulescu, D.F., Mihalca, A.D., Gherman, C.M., Leucuta, D., Meyer, F., Hizo-Teufel, C., Fingerle, V., Huber, I., 2016. Immunohistochemistry and real-time PCR as diagnostic tools for detection of *Borrelia burgdorferi* sensu lato in ticks collected from humans. Exp Appl Acarol. 69, 49–60. https://doi.org/10.1007/s10493-016-0012-y.

Camicas, J.L., 1998. The Ticks of the World (Acarida, Ixolida): Nomenclature, Described stages, Hosts, Distribution, Éditions de l’Orstom.

Courtney, J.W., Kostelnik, L.M., Zeidner, N.S., Massung, R.F., 2004. Multiplex real-time PCR for detection of *Anaplasma phagocytophilum* and *Borrelia burgdorferi*. J Clin Microbiol. 42, 3164–3168. https://doi.org/10.1128/JCM.42.7.3164-3168.2004.

Darling, A.E., Mau, B., Perna, N.T., 2010. progressiveMauve: multiple genome alignment with gene gain, loss and rearrangement. PLoS One. 5, e11147. https://doi.org/10.1371/journal.pone.0011147.

de Leeuw, B.H., Maraha, B., Hollemans, L., Sprong, H., Brandenburg, A.H., Westenend, P.J., Kusters, J.G., 2014. Evaluation of *Borrelia* real time PCR DNA targeting OspA, FlaB and 5S-23S IGS and *Borrelia* 16S rRNA RT-qPCR. J Microbiol Methods. 107, 41–46. https://doi.org/10.1016/j.mimet.2014.09.001.

Edgar, R.C., 2004. MUSCLE: multiple sequence alignment with high accuracy and high throughput. Nucleic Acids Res. 32, 1792–1797. https://doi.org/10.1093/nar/gkh340.

Estrada-Pena, A., Sprong, H., Cabezas-Cruz, A., de la Fuente, J., Ramo, A., Coipan, E.C., 2016. Nested coevolutionary networks shape the ecological relationships of ticks, hosts, and the Lyme disease bacteria of the *Borrelia burgdorferi* (s.l.) complex. Parasit Vectors. 9, 517. https://doi.org/10.1186/s13071-016-1803-z.

Ferdin, J., Cerar, T., Strle, F., Ruzic-Sabljic, E., 2010. Evaluation of real-time PCR targeting hbb gene for *Borrelia* species identification. J Microbiol Methods. 82, 115–119. https://doi.org/10.1016/j.mimet.2010.04.009.

Fraser, C.M., Casjens, S., Huang, W.M., Sutton, G.G., Clayton, R., Lathigra, R., White, O., Ketchum, K.A., Dodson, R., Hickey, E.K., Gwinn, M., Dougherty, B., Tomb, J.F., Fleischmann, R.D., Richardson, D., Peterson, J., Kerlavage, A.R., Quackenbush, J., Salzberg, S., Hanson, M., van Vugt, R., Palmer, N., Adams, M.D., Gocayne, J., Weidman, J., Utterback, T., Watthey, L., McDonald, L., Artiach, P., Bowman, C., Garland, S., Fuji, C., Cotton, M.D., Horst, K., Roberts, K., Hatch, B., Smith, H.O., Venter, J.C., 1997. Genomic sequence of a Lyme disease spirochaete, *Borrelia burgdorferi*. Nature. 390, 580–586. https://doi.org/10.1038/37551.

Gern, L., 2008. *Borrelia burgdorferi* sensu lato, the agent of lyme borreliosis: life in the wilds. Parasite. 15, 244–247. https://doi.org/10.1051/parasite/2008153244.

Gómez-Díaz, E., Doherty, P., Duneau, D., McCoy, K., 2010. Cryptic vector divergence masks vector-specific patterns of infection: an example from the marine cycle of Lyme borreliosis. Evolutionary Applications. 3, 391–401. https://doi.org/10.1111/j.1752-4571.2010.00127.x.

Gooskens, J., Templeton, K.E., Claas, E.C., van Dam, A.P., 2006. Evaluation of an internally controlled real-time PCR targeting the ospA gene for detection of *Borrelia burgdorferi* sensu lato DNA in cerebrospinal fluid. Clin Microbiol Infect. 12, 894–900. https://doi.org/10.1111/j.1469-0691.2006.01509.x.

Herrmann, C., Gern, L., Voordouw, M.J., 2013. Species co-occurrence patterns among Lyme borreliosis pathogens in the tick vector *Ixodes ricinus*. Appl Environ Microbiol. 79, 7273–7280. https://doi.org/10.1128/AEM.02158-13.

Ivanova, L.B., Tomova, A., Gonzalez-Acuna, D., Murua, R., Moreno, C.X., Hernandez, C., Cabello, J., Cabello, C., Daniels, T.J., Godfrey, H.P., Cabello, F.C., 2014. *Borrelia chilensis*, a new member of the *Borrelia burgdorferi* sensu lato complex that extends the range of this genospecies in the Southern Hemisphere. Environ Microbiol. 16, 1069–1080. https://doi.org/10.1111/1462-2920.12310.

Johnson, G., Nolan, T., Bustin, S.A., 2013. Real-time quantitative PCR, pathogen detection and MIQE. Methods Mol Biol. 943, 1–16. https://doi.org/10.1007/978-1-60327-353-4_1.

King, J.L., Smith, A.D., Mitchell, E.A., Allen, M.S., 2017. Validation of droplet digital PCR for the detection and absolute quantification of *Borrelia* DNA in *Ixodes scapularis* ticks. Parasitology. 144, 359–367. https://doi.org/10.1017/S0031182016001864.

Kralik, P., Ricchi, M., 2017. A Basic Guide to Real Time PCR in Microbial Diagnostics: Definitions, Parameters, and Everything. Front Microbiol. 8, 108. https://doi.org/10.3389/fmicb.2017.00108.

Lager, M., Faller, M., Wilhelmsson, P., Kjelland, V., Andreassen, A., Dargis, R., Quarsten, H., Dessau, R., Fingerle, V., Margos, G., Noraas, S., Ornstein, K., Petersson, A.C., Matussek, A., Lindgren, P.E., Henningsson, A.J., 2017. Molecular detection of *Borrelia burgdorferi* sensu lato - An analytical comparison of real-time PCR protocols from five different Scandinavian laboratories. PLoS One. 12, e0185434. https://doi.org/10.1371/journal.pone.0185434.

Larsson, A., 2014. AliView: a fast and lightweight alignment viewer and editor for large datasets. Bioinformatics. 30, 3276–3278. https://doi.org/10.1093/bioinformatics/btu531.

Leger, E., Liu, X., Masseglia, S., Noel, V., Vourc’h, G., Bonnet, S., McCoy, K.D., 2015. Reliability of molecular host-identification methods for ticks: an experimental in vitro study with *Ixodes ricinus*. Parasit Vectors. 8, 433. https://doi.org/10.1186/s13071-015-1043-7.

Machackova, M., Obornik, M., Kopecky, J., 2006. Effect of salivary gland extract from *Ixodes ricinus* ticks on the proliferation of *Borrelia burgdorferi* sensu stricto in vivo. Folia Parasitol (Praha). 53, 153–158. https://doi.org/10.14411/fp.2006.020.

Maes, L., Carolus, T., De Preter, V., Ignoul, S., Cartuyvels, R., Braeken, L., D’Huys, P.J., Saegeman, V., Kabamba, B., Raymaekers, M., 2017. Technical and clinical validation of three commercial real-time PCR kits for the diagnosis of neuroborreliosis in cerebrospinal fluid on three different real-time PCR platforms. Eur J Clin Microbiol Infect Dis. 36, 273–279. https://doi.org/10.1007/s10096-016-2797-3.

Mangold, A.J., Bargues, M.D., Mas-Coma, S., 1998. 18S rRNA gene sequences and phylogenetic relationships of European hard-tick species (Acari: Ixodidae). Parasitol Res. 84, 31–37. https://doi.org/10.1007/s004360050352.

Marconi, R.T., Garon, C.F., 1992. Development of polymerase chain reaction primer sets for diagnosis of Lyme disease and for species-specific identification of Lyme disease isolates by 16S rRNA signature nucleotide analysis. J Clin Microbiol. 30, 2830-2834. https://doi.org/10.1128/jcm.30.11.2830-2834.1992.

Margos, G., Fingerle, V., Reynolds, S., 2019. *Borrelia bavariensis*: Vector Switch, Niche Invasion, and Geographical Spread of a Tick-Borne Bacterial Parasite. FRONTIERS IN ECOLOGY AND EVOLUTION. 7. https://doi.org/10.3389/fevo.2019.00401.

Margos, G., Vollmer, S.A., Ogden, N.H., Fish, D., 2011. Population genetics, taxonomy, phylogeny and evolution of *Borrelia burgdorferi* sensu lato. Infect Genet Evol. 11, 1545–1563. https://doi.org/10.1016/j.meegid.2011.07.022.

Marsot, M., Chapuis, J.L., Gasqui, P., Dozieres, A., Masseglia, S., Pisanu, B., Ferquel, E., Vourc’h, G., 2013. Introduced Siberian Chipmunks (*Tamias sibiricus barberi*) Contribute More to Lyme Borreliosis Risk than Native Reservoir Rodents. PLoS One. 8, e55377. https://doi.org/10.1371/journal.pone.0055377.

Munoz-Leal, S., Ramirez, D.G., Luz, H.R., Faccini, J.L.H., Labruna, M.B., 2020. “*Candidatus* Borrelia ibitipoquensis,” a *Borrelia valaisiana*-Related Genospecies Characterized from *Ixodes paranaensis* in Brazil. Microb Ecol. 80, 682–689. https://doi.org/10.1007/s00248-020-01512-x.

Nunes, M., Parreira, R., Carreira, T., Inacio, J., Vieira, M.L., 2018. Development and evaluation of a two-step multiplex TaqMan real-time PCR assay for detection/quantification of different genospecies of *Borrelia burgdorferi* sensu lato. Ticks Tick Borne Dis. 9, 176–182. https://doi.org/10.1016/j.ttbdis.2017.09.001.

Nurk, S., Bankevich, A., Antipov, D., Gurevich, A., Korobeynikov, A., Lapidus, A., Prjibelsky, A., Pyshkin, A., Sirotkin, A., Sirotkin, Y., Stepanauskas, R., McLean, J., Lasken, R., Clingenpeel, S.R., Woyke, T., Tesler, G., Alekseyev, M.A., Pevzner, P.A., 2013. Assembling Genomes and Mini-metagenomes from Highly Chimeric Reads. Research in Computational Molecular Biology, 158–170. https://doi.org/10.1007/978-3-642-37195-0.

Ornstein, K., Barbour, A.G., 2006. A reverse transcriptase-polymerase chain reaction assay of *Borrelia burgdorferi* 16S rRNA for highly sensitive quantification of pathogen load in a vector. Vector Borne Zoonotic Dis. 6, 103–112. https://doi.org/10.1089/vbz.2006.6.103.

Portnoi, D., Sertour, N., Ferquel, E., Garnier, M., Baranton, G., Postic, D., 2006. A single-run, real-time PCR for detection and identification of *Borrelia burgdorferi* sensu lato species, based on the hbb gene sequence. FEMS Microbiol Lett. 259, 35–40. https://doi.org/10.1111/j.1574-6968.2006.00249.x.

Postic, D., Assous, M.V., Grimont, P.A., Baranton, G., 1994. Diversity of *Borrelia burgdorferi* sensu lato evidenced by restriction fragment length polymorphism of rrf (5S)-rrl (23S) intergenic spacer amplicons. Int J Syst Bacteriol. 44, 743–752. https://doi.org/10.1099/00207713-44-4-743.

Rauer, S., Kastenbauer, S., Hofmann, H., Fingerle, V., Huppertz, H.I., Hunfeld, K.P., Krause, A., Ruf, B., Dersch, R., Consensus, g., 2020. Guidelines for diagnosis and treatment in neurology - Lyme neuroborreliosis. Ger Med Sci. 18, Doc03. https://doi.org/10.3205/000279.

Rauter, C., Hartung, T., 2005. Prevalence of *Borrelia burgdorferi* sensu lato genospecies in *Ixodes ricinus* ticks in Europe: a metaanalysis. Appl Environ Microbiol. 71, 7203–7216. https://doi.org/10.1128/AEM.71.11.7203-7216.2005.

Schutzer, S.E., Body, B.A., Boyle, J., Branson, B.M., Dattwyler, R.J., Fikrig, E., Gerald, N.J., Gomes-Solecki, M., Kintrup, M., Ledizet, M., Levin, A.E., Lewinski, M., Liotta, L.A., Marques, A., Mead, P.S., Mongodin, E.F., Pillai, S., Rao, P., Robinson, W.H., Roth, K.M., Schriefer, M.E., Slezak, T., Snyder, J.L., Steere, A.C., Witkowski, J., Wong, S.J., Branda, J.A., 2019. Direct Diagnostic Tests for Lyme Disease. Clin Infect Dis. 68, 1052–1057. https://doi.org/10.1093/cid/ciy614.

Schwaiger, M., Peter, O., Cassinotti, P., 2001. Routine diagnosis of *Borrelia burgdorferi* (sensu lato) infections using a real-time PCR assay. Clin Microbiol Infect. 7, 461–469.

Stanek, G., Wormser, G.P., Gray, J., Strle, F., 2012. Lyme borreliosis. Lancet. 379, 461–473. https://doi.org/10.1016/S0140-6736(11)60103-7.

Strnad, M., Grubhoffer, L., Rego, R.O.M., 2020. Novel targets and strategies to combat borreliosis. Applied Microbiology and Biotechnology. 104, 1915–1925. https://doi.org/10.1007/s00253-020-10375-8.

Strube, C., Montenegro, V.M., Epe, C., Eckelt, E., Schnieder, T., 2010. Establishment of a minor groove binder-probe based quantitative real time PCR to detect *Borrelia burgdorferi* sensu lato and differentiation of *Borrelia spielmanii* by ospA-specific conventional PCR. Parasit Vectors. 3, 69. https://doi.org/10.1186/1756-3305-3-69.

Takano, A., Nakao, M., Masuzawa, T., Takada, N., Yano, Y., Ishiguro, F., Fujita, H., Ito, T., Ma, X., Oikawa, Y., Kawamori, F., Kumagai, K., Mikami, T., Hanaoka, N., Ando, S., Honda, N., Taylor, K., Tsubota, T., Konnai, S., Watanabe, H., Ohnishi, M., Kawabata, H., 2011. Multilocus sequence typing implicates rodents as the main reservoir host of human-pathogenic *Borrelia garinii* in Japan. J Clin Microbiol. 49, 2035–2039. https://doi.org/10.1128/JCM.02544-10.

Toohey-Kurth, K., Reising, M.M., Tallmadge, R.L., Goodman, L.B., Bai, J., Bolin, S.R., Pedersen, J.C., Bounpheng, M.A., Pogranichniy, R.M., Christopher-Hennings, J., Killian, M.L., Mulrooney, D.M., Maes, R., Singh, S., Crossley, B.M., 2020. Suggested guidelines for validation of real-time PCR assays in veterinary diagnostic laboratories. J Vet Diagn Invest. 32, 802–814. https://doi.org/10.1177/1040638720960829.

Tsao, J.I., Wootton, J.T., Bunikis, J., Luna, M.G., Fish, D., Barbour, A.G., 2004. An ecological approach to preventing human infection: vaccinating wild mouse reservoirs intervenes in the Lyme disease cycle. Proc Natl Acad Sci U S A. 101, 18159–18164. https://doi.org/10.1073/pnas.0405763102.

Venczel, R., Knoke, L., Pavlovic, M., Dzaferovic, E., Vaculova, T., Silaghi, C., Overzier, E., Konrad, R., Kolencik, S., Derdakova, M., Sing, A., Schaub, G.A., Margos, G., Fingerle, V., 2016. A novel duplex real-time PCR permits simultaneous detection and differentiation of *Borrelia miyamotoi* and *Borrelia burgdorferi* sensu lato. Infection. 44, 47–55. https://doi.org/10.1007/s15010-015-0820-8.

Vourc’h, G., Abrial, D., Bord, S., Jacquot, M., Masseglia, S., Poux, V., Pisanu, B., Bailly, X., Chapuis, J.L., 2016. Mapping human risk of infection with *Borrelia burgdorferi* sensu lato, the agent of Lyme borreliosis, in a periurban forest in France. Ticks Tick Borne Dis. 7, 644–652. https://doi.org/10.1016/j.ttbdis.2016.02.008.

Yang, Z., 1997. PAML: a program package for phylogenetic analysis by maximum likelihood. Comput Appl Biosci. 13, 555–556. https://doi.org/10.1093/bioinformatics/13.5.555.

Ye, J., Coulouris, G., Zaretskaya, I., Cutcutache, I., Rozen, S., Madden, T.L., 2012. Primer-BLAST: a tool to design target-specific primers for polymerase chain reaction. BMC Bioinformatics. 13, 134. https://doi.org/10.1186/1471-2105-13-134.

